# The direct regulation of *Aalbdsx* on *AalVgR* is indispensable for ovarian development in *Aedes albopictus*

**DOI:** 10.1101/2020.07.26.222224

**Authors:** Binbin Jin, Yijie Zhao, Peiwen Liu, Yan Sun, Xiaocong Li, Xin Zhang, Xiao-Guang Chen, Jinbao Gu

## Abstract

**BACKGROUND:** *Aedes albopictus* is an important vector with an extensive worldwide distribution. Only female mosquitoes play a significant role in the transmission of pathogens. *Doublesex* (*dsx*) is a central nexus gene in the insect somatic sex determination hierarchy.

**RESULTS:** In this study, we characterized the full-length sex-specific splicing forms of the *Ae. albopictus dsx* gene (*Aalbdsx*). Then, we identified 15 direct target genes of DSX in adult females using digital gene expression (DGE) combined with qPCR following a chromatin immunoprecipitation assay (ChIP) with specific DSX antibodies. The knockdown of *Aalbdsx* suppressed ovarian development, and the transcript levels of the *Aalbdsx* target *vitellogenin receptor* (*VgR*) gene decreased, whereas *vitellogenin* (*Vg*) expression showed an increase in the fat body. Genes in the major *Vg* regulatory pathway were also upregulated. Our results suggest that the effects of *Aalbdsx* RNAi on ovarian development are exerted mainly via *VgR* rather than *Vg*.

**CONCLUSION:** The results of our study not only provide a reference for the further elucidation of the sex determination cascade and comparative analyses of *dsx* target interactions in mosquitoes but also reveal potential molecular targets for application to the development of sterile male mosquitoes to be released for vector control.

## 1. Introduction

Sex determination in most insects is controlled by a complex hierarchical regulatory cascade. Once a primary signal at the top of the cascade is triggered, it is transmitted through a cascade of sex determination genes to a downstream master-switch gene, that regulates somatic sexual differentiation in both sexes. The primary signals for sex determination are highly diverse in different species. For example, the dosage of the X chromosome is the primary signal for *Drosophila* sex determination(1). In other dipterans, male-determining factors (M-factors) such as *Musca domestica male determiner* (*Mdmd*)(2), *Maleness-on-the-Y* (*MoY*)(3) and *Nix*(4, 5), are employed to determine male sex in *Musca domestica, Ceratitis capitata* (*Medfly*), *Aedes aegypti and Ae. albopictus*, respectively. In lepidopteran insects, a female-specific piRNA produced from a piRNA precursor feminizing gene (*Fem*) arranged in tandem in the sex-determining region of the W chromosome, is the top signal in the *Bombyx mori* sex determination cascade(6). In the hymenoptera insect *Apis mellifera*, the initial sex-determining signal is provided by heterozygosity at the *complementary sex determiner* (*csd*) locus(7). In the coleopteran insect *Tribolium castaneum*, the maternally deposited *transformer* (*tra*) transcript is necessary for the initiation of the positive autoregulatory feedback loop in females, which guides the female-specific developmental program(8). The downstream genes in the hierarchical regulatory cascade are generally more conserved than the upstream genes, such as the *Drosophila melanogaster doublesex (dsx)* gene, which acts as the terminal “double-switch” in the sex determination pathway. Orthologs of *dsx* have been identified in almost all insect species investigated thus far, covering a range of Phthiraptera(9), Hemiptera(9), Diptera(10-15), Lepidoptera(16, 17), Coleoptera(18, 19), Strepsiptera(9) and Hymenoptera(20-22), leading to the hypothesis that the sex determination cascade evolves from the bottom up(23).

DSX proteins are members of the doublesex and male abnormal-3-related transcription factor (Dmrt) family, which are structurally and functionally conserved and are classified in a group of zinc-finger (ZnF) proteins with dimeric sequence-specific DNA-binding motifs that regulate transcription and play a key role in sex determination and/or differentiation throughout different phyla(24-26). In insects, the *dsx* gene is constitutively transcribed during larval development and in the soma and germlines of adults of both sexes; however, its precursor messenger RNA (pre-mRNA) is subjected to sex-specific alternative splicing (AS) regulation so that distinct male- and female-specific transcripts are produced in each sex, which are translated into identical male-specific DSX^M^ and female-specific DSX^F^ proteins. It has been established that the DSX proteins and the *fruitless* (*fru*) product FRU protein control nearly all aspects of somatic sexual differentiation, including anatomical and behavioral differences. However, the target genes of DSX have only been predicted and analyzed on a genome-wide scale in *D. melanogaster*(27-29) and in beetles (*Tribolium castaneum and Onthophagus taurus*)(19, 30). Furthermore, only a few genes, such as the *Vg*(13, 17, 19, 29, 31-35), *pheromone binding protein* (*pbp*) gene(36, 37), *hexamerin*(17, 36), *intersex* (*ix*)(38), *bric-a-bric* (*bab*)(28, 29, 39, 40), *fatty acid desaturation 2* (*fad2*)(41), and *wingless* (*wg*)(42) genes have been functionally demonstrated to be direct DSX targets in previous studies.

Among mosquitoes, *Drosophila dsx* orthologs were previously identified *in Ae. aegypti*(14), *Anopheles gambiae*(10), *and Culex pipiens*(43), and sex-specific splicing and conserved domains are maintained in these orthologs. The RNA interference (RNAi)-mediated silencing of *dsx*^F^ expression in the larvae of *A. aegypti* by the feeding of double-stranded RNAs leads to an increase in the mortality rate of female larvae, resulting in a highly male-biased population of mosquitoes(44). Furthermore, the downregulation of *dsx* in *Ae. aegypti* pupae by siRNA microinjection disrupts multiple sex-specific morphological characteristics, causing significant decreases in adult female wing size, the length of the antenna and maxillary palps, ovary length, ovariole length, ovariole number, and even the female lifespan(45). However, there is still a lack of publicly available data elucidating the major DSX target genes in mosquitoes.

The Asian tiger mosquito, *Ae. albopictus* (Diptera: Culicidae), a mosquito native to Asia, shows a rapidly increasing global range and is ranked as one of the 100 most successful invasive species in the world(46, 47). *Ae. albopictus* has been showen to be a competent vector that can experimentally transmit at least 26 arboviruses, including the dengue (DENV), chikungunya (CHIKV), yellow fever (YFV) and Zika (ZIKV) viruses(48). Since only female mosquitoes play a key role in the transmission of pathogens, we isolated and characterized full-length *dsx* cDNAs from *Ae. albopictus* and focused on the DSX target gene in female adults. RNA interference (RNAi) was used to target the common region of female-specific *dsx* isoforms in female adults, and digital gene expression (DGE) analysis was performed using the Illumina NGS platform to screen for genes showing significant changes in expression levels in response to *Aalbdsx*^F^ downregulation. Then, the direct target genes of the DSX protein were confirmed by the qPCR method using immunoprecipitated DNA as a template following a chromatin immunoprecipitating assay (ChIP) with specific DSX antibodies. Finally, several direct *dsx* target genes in *Ae. albopictus* adult females were confirmed. The effects of *dsx* repression on ovarian development were investigated, and the response of the major signaling pathway in the regulation of *Vg* post-*dsx* RNAi was further analyzed. We found that the effects of *Aalbdsx* RNAi on ovarian development were exerted mainly via *VgR* rather than *Vg*. The data presented here confirm the evolutionarily conserved role of *dsx* in insect sexual differentiation.

## 2. Materials and methods

### 2.1 Mosquitoes

The *Ae. albopictus* Foshan strain was a kind gift from the Centers for Disease Control of Guangdong Province. The strain was isolated from Foshan, Guangdong, P.R.C and has been established in the laboratory since 1981. All mosquitoes were reared in a climate-controlled insectary at 27 ± 1°C with a relative humidity of 70%-80% under a 14 L:10D photoperiod. The larvae were reared in pans and fed small turtle food (INCH-GOLD, Shenzhen, China). Adult mosquitoes were kept in 20 cm*30 cm*45 cm yarn cages and allowed access to a cotton wick soaked in 10% glucose as a carbohydrate source. Adult females were allowed to take blood meals from defibrinated sheep blood (Solarbio Life Sciences, Beijing, China) supplied through glass membrane feeders covered by a porcine membrane(49).

### 2.2 *Aalbdsx* cloning strategy

tBLASTn searches of the *Ae. albopictus* transcripts and genome (http://www.vectorbase.org/) were performed using *Ae. aegypti* female-specific DSX (Aeadsx^F^, GenBank: ACY78700) (14) as a query. To determine the molecular organization of *Aalbdsx* following RT-PCR using sequence-specific primers, 3’- and 5’-RACE analyses were performed to determine the transcription start sites and the 3’ ends of the genes. 5’-RACE and 3’-RACE reactions were performed with the Smart RACE cDNA Amplification Kit (Clontech, Palo Alto, CA) according to the manufacturer’s instructions. The sequences of the primers used in the RT-PCR and RACE analyses for the amplification of nearly full-length cDNAs are shown in Table S2. The amplified cDNA fragments were purified in agarose gels, eluted, subcloned into the pJET1.2/blunt vector (Thermo Fisher Scientific, Carlsbad, CA) and confirmed by sequencing (Shanghai Invitrogen Biotechnology Co, Ltd.).

### 2.3 In Vivo RNA Interference

*Aalbdsx*-specific PCR primers were synthesized with the T7 RNA polymerase promoter sequence appended to their 5′ends. cDNA was used as a template to amplify fragments of *Aalbdsx*. The purified PCR products were used as templates to synthesize dsRNA using the T7 RiboMAX Express RNAi System (Promega Corporation, Madison WI, USA) according to the manufacturer’s instructions. Green fluorescent protein (GFP) dsRNA was used as a control. The primers used for amplifying the templates of *Aalbdsx* and GFP mRNA are listed in Table S3.

Two-day-old *Ae. albopictus* female adults were anesthetized on ice, and approximately 500 nL of dsRNA solution (2000 ng/µl) was then injected into their thorax under a microscope, as described previously(50). After injection, the adult mosquitoes were immediately transferred to small plastic cups (900 ml, 11 cm diameter at the top) and fed with a 10% glucose solution through soaked cotton wicks and allowed to recover. Three independent biological replicates were included for each treatment (n = 30 per replicate), followed by digital gene expression (DGE) profiling analysis. The knockdown efficiency of *Aalbdsx* expression in the injected mosquitoes was confirmed by qRT-PCR with gene-specific primers, and *Ae. albopictus ribosomal protein S7* (*AalrpS7*, GenBank: JN132168.1) gene was used as an endogenous control. The knockdown efficiencies of all primers used in these experiments are listed in Table S3.

### 2.4 RNA extraction, DGE Library Preparation and Sequencing

Total RNA was isolated from each sample using the TRIzol reagent according to the manufacturer’s instructions (Invitrogen, Carlsbad, CA). RNA purity was examined with a Nano Photometer spectrophotometer (IMPLEN, CA). The RNA concentration was measured with a Qubit® RNA Assay Kit in a Qubit 2.0 Florometer. RNA integrity was assessed using the RNA Nano 6000 Assay Kit on a Bioanalyzer 2100 system (Agilent Technologies, CA). Two samples with high RNA integrity numbers (RINs) from each treatment were selected for library preparation. Briefly, the total RNA samples were treated with RNase-free deoxyribonuclease I (DNase I) (New England BioLabs, Beverly, MA) to remove residual DNA. Then, the total RNA was treated with Oligo (dT) magnetic beads to capture mRNA. Next, the purified mRNA was fragmented using fragmentation buffer. The first strand cDNA was reverse transcribed using SuperScript III Reverse Transcriptase (Invitrogen, Carlsbad, CA) with random primers. The second strand of cDNA was synthesized using DNA polymerase (New England BioLabs, Beverly, MA) and then end repaired, followed by 3’single nucleotide A (adenine) addition and the ligation of adapters. After ligation with sequencing adapters, PCR amplification was performed, and sequencing was conducted on an Illumina HiSeq 2000 sequencer at the Beijing Genomics Institute (BGI, Shenzhen, China). The sequencing data have been deposited at the NCBI Gene Expression Omnibus (GEO) database/NCBI Sequence Read Archive (SRA) repository (accession Nos. SRR10290622, SRR10290623, SRR10290624 and SRR10290625).

### 2.5 Analysis of DGE Tags

After the removal of low-quality reads, adaptors and reads containing poly-N sequences, the clean reads were filtered from the raw reads, and the clean data were aligned to the reference genome sequence of the *Ae. albopictus* Foshan strain (accession number is SRA215477)(51) using the program TopHat version 2.0.9(52). The default settings for the tolerance parameters were used (no more than two mismatches). The read numbers mapped to each gene were counted, and expression abundance was calculated based on the number of reads per kilobase of transcript per million mapped reads (RPKM). Differential expression analysis of two samples was performed in R using the DESeq2 Bioconductor package(53). Thresholds of a false discovery rate (FDR) <0.005 and absolute value of the log 2-fold-change > 1 were used to determine significant differences in gene expression(54).

### 2.6 Quantitative real-time PCR validation of RNA-Seq data

Fourteen candidate genes were selected for validation using quantitative real-time PCR (RT-qPCR). The candidate genes and their primers employed for RT-qPCR are listed in Table S4. All RT-qPCR assays were performed in an ABI 7500 System with the SuperReal PreMix Plus kit (SYBR Green) (Tiangen, Beijing, China). All RT-qPCR procedures for each gene were performed in three biological replicates, with three technical repeats per experiment. Relative gene expression was normalized to the internal control gene *rpS7* and analyzed using the 2^−ΔΔCT^ method(55). The data are presented as the mean ± SD (n = 3). Significant differences among the experimental groups were analyzed in GraphPad Prism 8 using the unpaired t-test. The P value threshold was set at 0.05 (p < 0.05) for highly significant differences (*).

### 2.7 Chromatin immunoprecipitation (ChIP-qPCR) analysis

The physical binding of DSX to the direct target gene promoter 2 days post-emergence in adult females was studied by ChIP-qPCR analysis. The ChIP assay was performed using the SimpleChIP® Plus Sonication Chromatin IP Kit (Cell Signaling Technology, Beverly, MA) following the manufacturer’s protocol with some modifications(56). Sonication was performed under the following conditions: ultrasonic cell pulverizer (JY92-II, Shanghai Xinzhi, China) at a power of 15, sonication for 10 sec and resting on ice for 40 sec, repeated 15 times. After sonication, 2% of the sample was left as the input group, and the rest was reacted with IgG and *Aalbdsx* antibodies (2 µg, Convenience Biology, ChangZhou, China). Then, the immunoprecipitated DNA fragments were quantified by qPCR. The primers used for ChIP-qPCR are listed in Table S5. The ChIP-qPCR values obtained for the immunoprecipitated samples were normalized to the percentages of the respective inputs from 3 independent experiments. In brief, the amount of genomic DNA coprecipitated with anti-Aalbdsx or anti-IgG antibodies was calculated as a percent of the total input as follows: ΔCt [normalized ChIP] = (Ct [ChIP] - (Ct [Input] -Log_2_ (Input Dilution Factor))); % Input = 2 ^(-ΔCt [normalized ChIP])^.

### 2.8 Ovarian measurements and fat body dissection

To explore the effect of *Aalbdsx* RNAi on the ovarian development of *Ae. albopictus* adult females, 24 h post-emergence adult females were coupled with males and then injected with dsRNA, followed by routine rearing. At 36 h post-injection (p.i.), the mosquitoes were fed defibrinated sheep blood, and only freshly blood-fed female mosquitoes with obviously engorged, bright red abdomens were selected for subsequent analysis. At 48 h post-blood meal (PBM), the ovaries were dissected by tearing off the soft cuticle between the fifth and sixth abdominal sternites with a fine needle and then pulling off and placing the terminal segments in a drop of suitable mosquito saline buffer(57). The ovaries were dissected, and two measurement indicators were used to evaluate ovarian development: (1) the numbers of developed follicles and (2) individual follicle size.

To explore the effect of *Aalbds*x RNAi on the transcription levels of *AalVg* and *AalVgR* in the fat body and ovary post-blood meal (PBM), 24 h postemergence adult females were coupled, injected, and fed according to the procedure described above. The ovary and fat body tissues were then dissected from surviving female mosquitoes (48 h PBM) according to the protocol described previously(58). However, only the fat bodies and ovaries of individuals with obvious ovarian defects (left-right ovarian asymmetry, undeveloped ovaries, etc.) were collected for *AalVg* analysis. (ovarian defects=47, total =77).

### 2.9 qPCR analysis of injected mosquitoes

The inhibitory effect of *Aalbdsx* and the relative expression levels of *AalVgs, AalVgR* (AALF004868(59)) and the genes in the *dsx* regulation pathway (Table S6) were quantified by real-time PCR using the SuperReal PreMix Plus kit (SYBR Green) according to the manufacturer’s instructions. Total RNA was extracted from whole adult (non-blood-fed), fatbody and ovary (blood-fed) female mosquitoes in the RNAi and control groups (84 h p.i., 48 h PBM), using TRIzol according to the manufacturer’s directions.

### 2.10 Statistical Analysis

All experiments were carried out in triplicate. Data are expressed as the mean ± SD, and the statistical analysis of in vivo RNA interference, the relative expression levels of corresponding genes and egg counts were evaluated using unpaired Student’s test. Standard one-way ANOVA was used to evaluate the average follicle size of ovaries. No statistical methods were used to predetermine the sample size. The experiments were not randomized and the investigators were not blinded to allocation during the experiments and outcome assessment. Statistical significance was assigned when p values were <0.05 according to GraphPad Prism Version 8.02 and IBM SPSS Statistics 20.0.

## 3. Results

### 3.1 Isolation and molecular characterization of the *Aalbdsx* gene

A tBLASTn search of the *Ae*.*albopictus* Foshan strain transcripts and genome using *Aeadsx*^F^ as a query was performed to identify the putative ortholog of the *Aeadsx* gene. At an e-value of less than 1e^-10^, draft sequences (AALF017777) located on the following scaffolds were identified: JXUM01S004167 and JXUM01S006384.

The molecular cloning of *Aalbdsx* was performed by combining traditional PCR with 3’ and 5’ RACE analyses using the adult female and male cDNAs as templates. The sex-specific splicing transcripts of *Aalbdsx*, including three female-specific isoforms and one male-specific transcript, were identified and deposited in GenBank (*Aalbdsx*^*F1*^ (GenBank: MT218330), *Aalbdsx*^*F2*^ (GenBank: MT218331), *Aalbdsx*^*F3*^ (GenBank: MT218332), and *Aalbdsx*^*M*^ (GenBank: MT218333). The alignment of the *Aalbdsx* cDNA sequences with the corresponding genomic sequence was performed to define the exon/intron organization and AS events (Fig.1A). Fig.1A shows the molecular organization of the *Aalbdsx* RNA molecules. *Aalbdsx*^*F1*^ had a length of 4,237 bp and was composed of nine exons and eight introns. Exons 6 and 7 were alternatively spliced female-specific exons. *Aalbdsx*^*F1*^ encodes an ORF of 281 amino acids, beginning with an AUG located within exon 2, and the transcription start site is located at –1,510 bp from the initial AUG of the ORF. Compared with *Aalbdsx*^*F1*^, *Aalbdsx*^*F2*^ displayed an AS event of mutually exclusive exons, in which female-specific exon 6 was excised, whereas exon 7 was retained in the mRNAs after splicing. As a result, the termination codon was located in exon 7, and the predicted ORFs of *Aalbdsx*^*F2*^ could theoretically encode a protein of 270 amino acids. The *Aalbdsx*^*F3*^ mRNA splicing pattern differed from that of *Aalbdsx*^*F1*^ in that intron 6 was retained, and this intron retention event connected exon 6 and exon 7 to form a single continuous exon; however, *Aalbdsx*^*F3*^ exhibited the same ORF as *Aalbdsx*^*F1*^. Male-specific *Aalbdsx*^*M*^ consisted of 3,354 bp and encoded an ORF of 506 amino acids, using the same translation initiation codon as *Aalbdsx*^*F1*^, but lacked female-specific exons 6 and 7.

**Figure 1.**
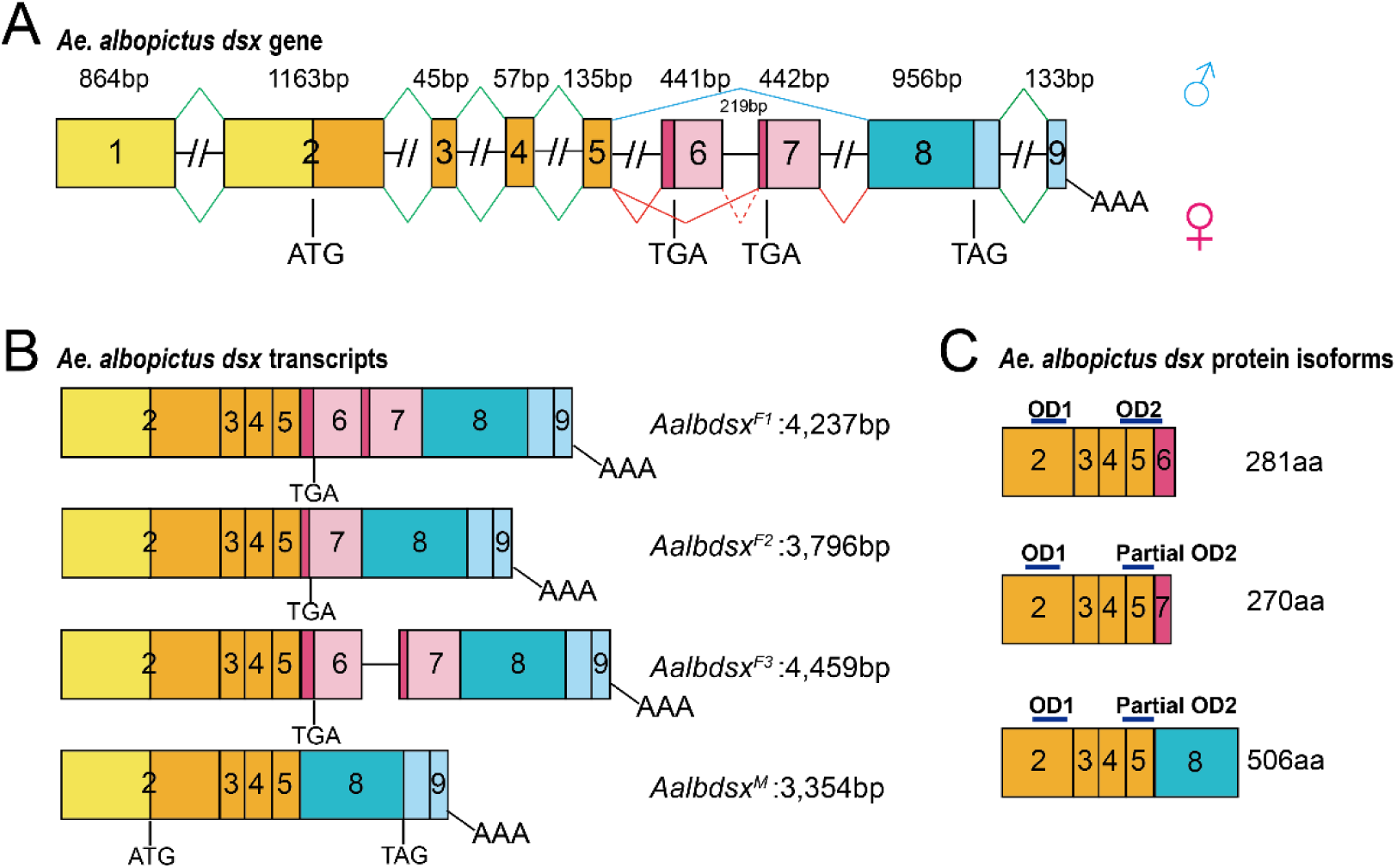
Molecular organization of *dsx* in *Ae. albopictus*. (A) Genomic organization of *Aalbdsx*. Female-specific and Male-specific exons/protein are indicated in pink and blue, respectively. Red dotted line within intron 6 represents a 219-bp intronic sequence alternatively removed in *Aalbdsx* transcripts of female. Exons and introns are not shown to scale. Dotted lines for introns indicate that their lengths remain unknown.1-9 is the exon numbering. Start (ATG) and stop (TGA/TAG) codons and poly (A) addition sites (AAA) are marked. (B) Splicing variants of *Ae. albopictus*. (C) Overview of *Ae. albopictus* male and female DSX isoform proteins, including the functional protein domains. The first oligomerization domain(OD1) is located at the N-terminus, whereas the second oligomerization domain (OD2) is located towards the C-terminus.

### 3.2 DGE library sequencing and analysis and qRT-PCR validation of differentially expressed genes

Four DGE libraries (two biological replicates were performed for each treatment) were produced from the whole body of *Aalbdsx*-RNAi adult females and *gfp*-dsRNA-injected adult females (negative control) using Illumina Solexa sequencing technology. As a result, a total of 12,744,233 and 14,878,358 raw tags were generated from the *Aalbdsx*-RNAi DGE libraries, 14,615,115 and 14,180,772 raw tag were generated from the control libraries. Then, a total of 11,570,176, 13,488,108, 13,299,143 and 12,862,091 clean tags, corresponding to 6,631,951, 7,623,850, 7,591,689 and 7,336,850 distinct clean tags, were filtered from the raw tags. All data are available in the NCBI Sequence Read Archive (SRA) under accession numbers SRR10290622, SRR10290623, SRR10290624 and SRR10290625.

To compare differentially expressed genes between the libraries, the level of gene expression was determined by normalizing the number of unambiguous tags in each library to RPKM values, and unigenes were considered significant at an FDR <0.005 and |log 2 fold_change| > 1. In comparison with the control, the expression levels of 60 unigenes were significantly different in *Aalbdsx*-knockdown adults (Table S1), including 33 upregulated genes and 25 downregulated genes. Intriguingly, an *Ae. Albopictus*, the *vitellogenin-A1 precursor* gene (AALF008766) was included in the list, while seven of the genes encoded a putative salivary secreted protein (AALF010748) or predicted salivary secreted peptides/proteins (AALF004423, AALF028226, AALF004632, AALF004765, AALF023211, AALF018059), and two genes (AALF018601 and AALF015062) encoded odorant-binding protein (OBP) or predicted OBPs. Among these genes, a gene with a |log 2 fold_change| greater than 2 was selected for further qRT-PCR analysis to verify the reliability of the DGE results in the present study. As a result, the relative expression levels of 10 upregulated and 4 downregulated genes were assessed. As shown in Fig.2, the *Aalbdsx* mRNA transcript level in the *Aalbdsx* knockdown groups was successfully depleted to 11.16 ± 3.01% (p<0.05) of the level in the *gfp* interference negative control group. Significant changes in *AalVg* expression were observed between the *Aalbdsx*-RNAi group and the *gfp*-RNAi control. Four genes were significantly downregulated in the *Aalbdsx*-RNAi group, whereas ten genes were significantly upregulated, showing the same expression pattern as in the initial DGE analysis (Table S1). These results confirmed the reliability of the DGE profiling analysis.

**Figure 2.**
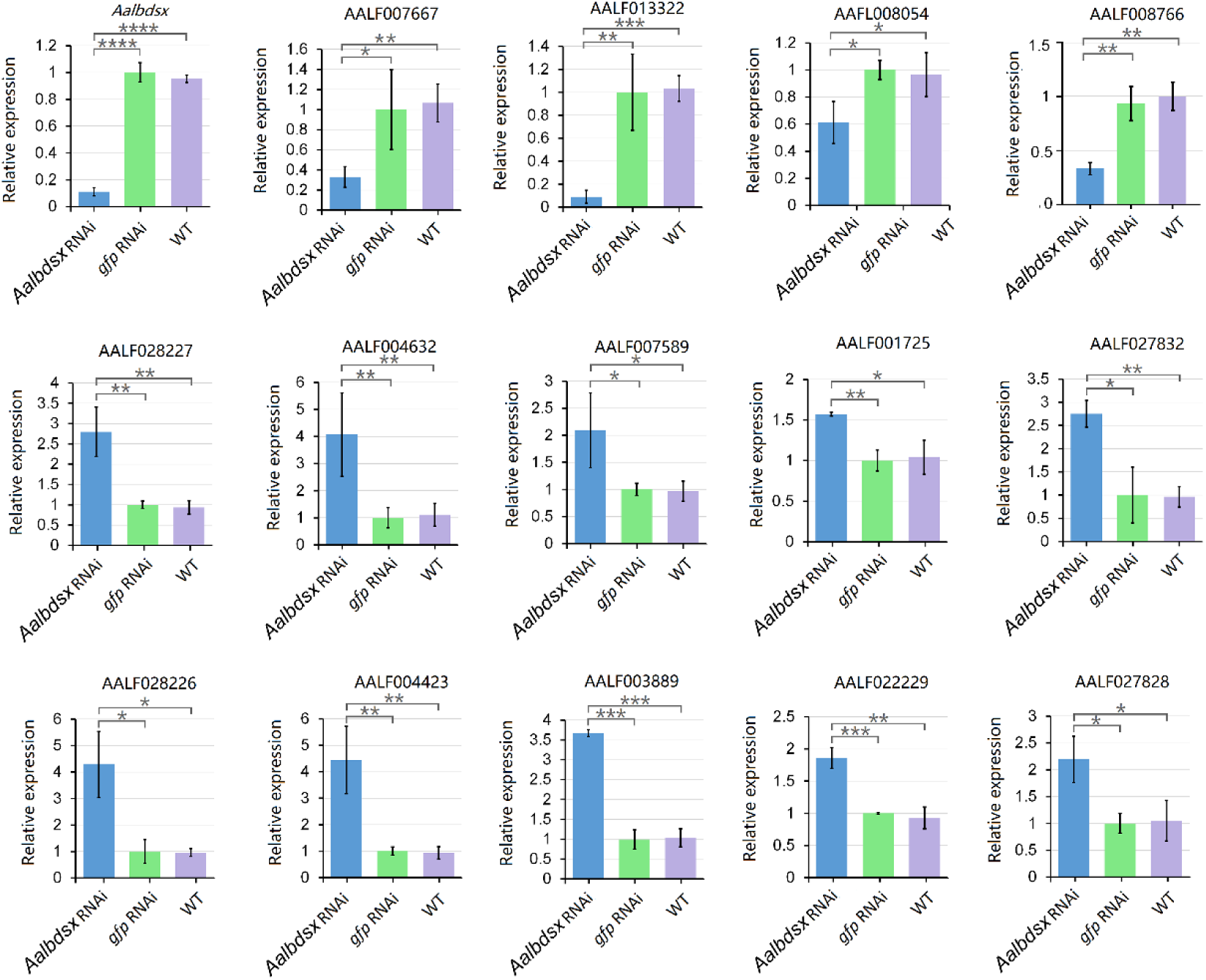
Validation of DGE expression profiles by qRT-PCR. *Aalbdsx* mRNA transcript levels were significantly repressed in the *Aalbdsx* RNAi groups compared to the control. DGE expression profiles of 14 differentially expressed genes (|log 2 ratio| ≥ 2) between the *Aalbdsx* RNAi group and the negative control group were confirmed by qRT-PCR. *Ae. albopictus ribosomal protein 7* (*rpS7*) mRNA was used as an internal control. Error bars indicate standard deviation. Data represent the mean ± SD (n = 3, *p < 0.05, **p < 0.01, and ***p < 0.001).

### 3.3 Chromatin immunoprecipitation (ChIP) qPCR analysis

ChIP-qPCR assays were performed to quantify the amount of the promoter DNA of candidate target genes precipitated by anti-Aalbdsx antibodies. The selected candidate target genes were categorized into three groups: the first group included the top 5 upregulated genes (AALF027828; AALF022229; AALF003889; AALF004423 and AALF028226) and the top 6 downregulated genes (AALF015200; AALF018059; AALF008054; AALF013322; AALF007667 and AALF008766) identified in *Aalbdsx* RNAi mosquitoes via DGE; the second group contained 4 orthologs of previously confirmed direct targets of the *Vg*-related gene, including three *Vg*-orthologous genes and one *VgR* gene from the *Ae. albopictus* Foshan strain *AalVg-A1* (AALF008766), *AalVg-B* (AALF019930), *AalVg-C1* (AALF008379), *AalVg-C2* (AALF027368) and *AalVgR* (AALF004868) (AALF008766 overlapping with group I); and the third group consisted of 7 putative or predicted salivary secreted protein encoding genes (AALF010748 AALF004423, AALF028226, AALF004632, AALF004765, AALF023211, and AALF018059, among which AALF004423, AALF028226, and AALF018059 overlapped with group I). As shown in Fig.3, a total of 15 genes presented significant binding of endogenous *Aalbdsx* to their promoter region in an analysis with gene-specific primers. Taken together with results of the DGE and qPCR analyses (Fig.2), these assays indicate that these genes are direct transcriptional targets of endogenous *Aalbdsx*.

**Figure 3.**
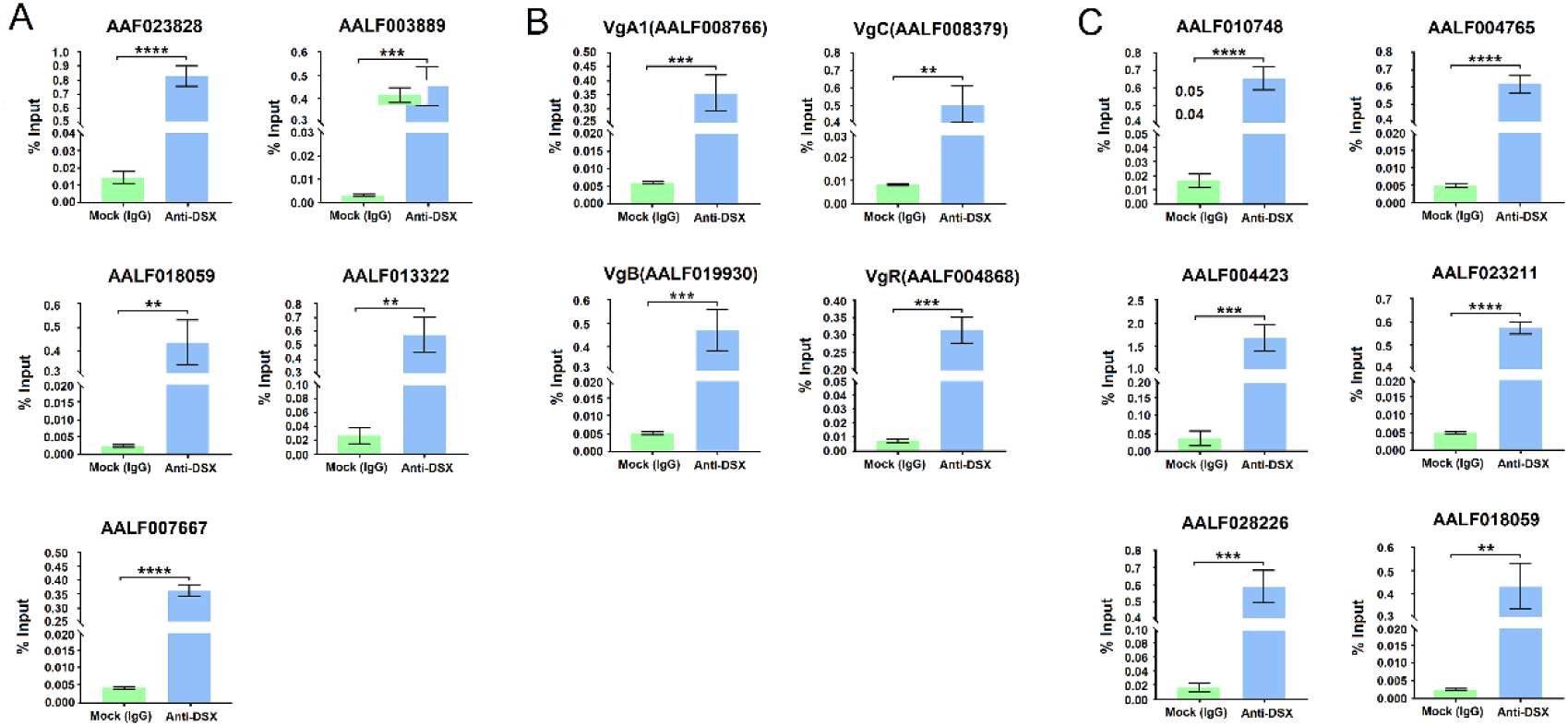
ChIP-qPCR analysis of candidate target genes of *Aalbdsx*. ChIP-qPCR assays were used to quantify the amount of promoter DNA of candidate target genes precipitated by anti-Aalbdsx antibodies, and the results were graphed as % input. Data are shown as the mean ± SD (unpaired t-test, n = 3, *p < 0.05, **p < 0.01, and ***p < 0.001). Error bars indicate standard deviation. (A) Top eleven differentially expressed genes in *Aalbdsx* RNAi mosquitoes compared with the control group, identified via DGE. (B) The orthologs of previously confirmed direct targets of *Vg*-related genes. (C) The differentially expressed genes encoding putative or predicted salivary secreted proteins.

### 3.4 Knockdown of *Aalbdsx* suppresses ovarian development

*Vgs* are major yolk protein precursors (YPPs), that play a key role in mosquito ovarian development. *Vgs* are primarily synthesized in the fat body and secreted into the hemolymph, then taken up by the ovaries via endocytosis mediated by the corresponding receptor(60). Although *Vg* is thought of as a relatively conserved target gene of *dsx* in insects(19), our DGE library sequencing and ChIP-qPCR results confirmed that *Vg* is also a target gene of *dsx* in *Ae. Albopicuts*. We investigated the effect of the RNAi knockdown of the *Aalbdsx* genes on ovarian development of adult females PBM. The mosquitoes in each group were dissected 48 h p.b.m (84 hours post-injection), and the knockdown efficiency of *Aalbdsx* was assessed by RT-qPCR. *gfp*-dsRNA-injected mosquitoes were used as a negative control, and wild-type mosquitoes were used as a blank control.

The average length of the follicles was measured, and the number of ovarioles in each individual was counted. As shown in Fig.4A, *Aalbdsx* transcript levels were successfully suppressed to 19.07 ± 8.72% of the levels in the control group (p<0.05), at 48 h.p.m. As expected, *Aalbdsx* downregulation led to a significant reduction (p<0.05) in the number of ovarioles in each individual (38.06 ± 15.72, n=47) compared with the negative control treatment (52.70 ± 13.40, n=37) and the untreated wild-type group (51.73 ± 4.12, n=37) (Fig.4C). Furthermore, some individuals showed obvious growth arrest and even a lack of ovarian development (Fig.4B). There were no statistically significant differences between the *gfp*-dsRNA injection group and wild-type group with regard to the ovariole number or average follicle size.

**Figure 4.**
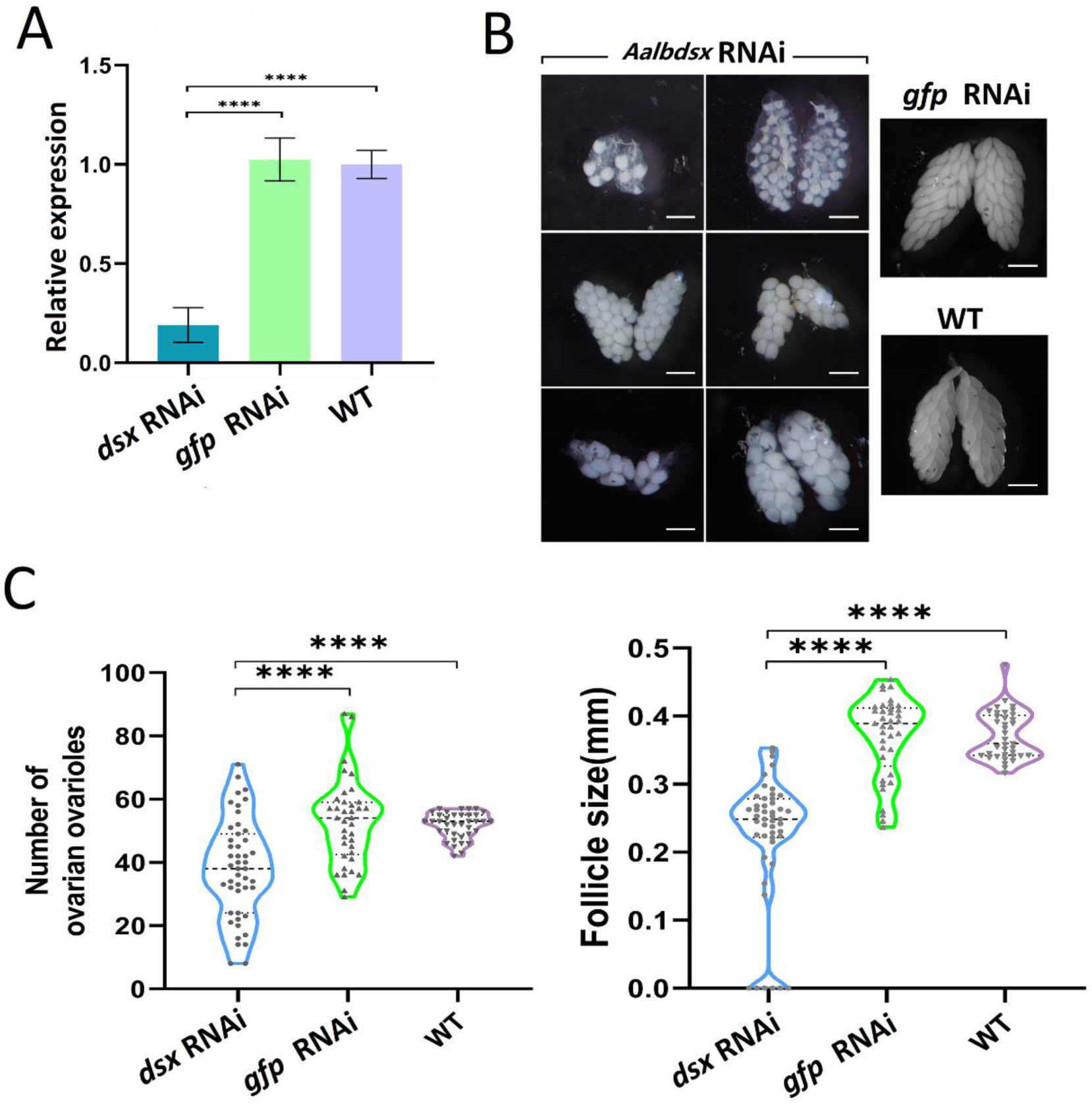
Effect of *Aalbdsx* knockdown on ovarian development in adult female *Ae. albopictus*. (A) Relative expression of *Aalbdsx* in *Aalbdsx* dsRNA injected groups compared with the *gfp* dsRNA-injected control and WT groups. The x-axis indicates the groups; WT indicates wild-type mosquitoes that did not receive any treatment. The expression of the *Aalbdsx* gene in the WT group was set as 1 and used to normalize *Aalbdsx* expression. The error bars represent the SD (n = 3, *p < 0.05, **p < 0.01, and ***p < 0.001). (B) Representative images of ovaries dissected from WT mosquitoes and mosquitoes from the *Aalbdsx* dsRNA-injected groups and *gfp* dsRNA-injected groups at 48 h PBM. The images were taken under a light microscope with a Nikon SMZ1000 stereomicroscope. (C) Number of ovarian ovarioles per female in the *Aalbdsx* dsRNA-injected groups compared with the *gfp* dsRNA-injected control and WT groups. Average follicle size (length of the long axis) in ovaries isolated from female mosquitoes in *Aalbdsx* dsRNA-injected groups compared with *gfp* dsRNA-injected control and WT groups. Scale bar: 500 μm.

### 3.5 The effects of *Aalbdsx* RNAi on ovarian development are mediated mainly via *AalVgR*, rather than *AalVg*

To determine whether the suppression of ovarian development in *Aalbdsx* RNAi mosquitoes mainly results from the influence of the expression of the *Aalbdsx* target gene *Vg*, the relative transcript levels of four *Vg*-orthologous genes in *Ae. albopictus* Foshan strain *AalVg-A1* (AALF008766), *AalVg-B* (AALF019930), *AalVg-C1*(AALF008379) and *AalVg-C2*(AALF027368) were compared between the *Aalbdsx* RNAi groups and the control groups at 48 h PBM (because of the high nucleotide sequence identity between *AalVg-C1* and *AalVg-C2*, the sum of the transcript levels of the two genes was quantitatively analyzed). The *Aalbdsx* transcript level was shown to be efficiently repressed in the *Aalbdsx* knockdown groups (Fig.5A). Interestingly, however, the mRNA transcript levels of all the *AalVg* genes, including *AalVg-A1, AalVg-B*, and *AalVg-C1/C2*, were significantly higher in the *Aalbdsx* knockdown groups than in the *gfp* dsRNA-injected groups (by up to 854.88 ± 269.15%, 4126.49 ± 1166.17% and 2492.22 ± 1217.89%, respectively, in the fat body and 270.49 ±74.85%, 889.31 ±445.82%, 984.51 ±167.20% in the ovary, p < 0.05). No significant changes in *AalVg* expression were observed between the *gfp* dsRNA-injected groups and the WT (Fig.5A). The upregulation of *Vg* was apparently contrary to the suppression of ovarian development. *Vg* is taken up in the developing ovaries via endocytosis by its receptor, *VgR. VgR* gene were identified as targets of *TcDsx*(19), and *AalVgR* was verified to be a target gene of *Aalbdsx* by ChIP-qPCR. Therefore, we further examined the relative *AalVgR* transcript levels in adults with *Aalbdsx* downregulation at 48 h PBM As expected, *AalVgR* transcript levels were significantly repressed in the *Aalbdsx* knockdown groups to 2.41 ± 0.71% and 42.35 ± 6.17% of the levels in the *gfp* dsRNA-injected groups in the fat body and ovary, respectively (p < 0.05). These results indicate that the suppression of *AalVgR* leads to a disorder of the ovarian uptake of *AalVg* from the hemolymph and ultimately blocks ovarian development.

**Figure 5.**
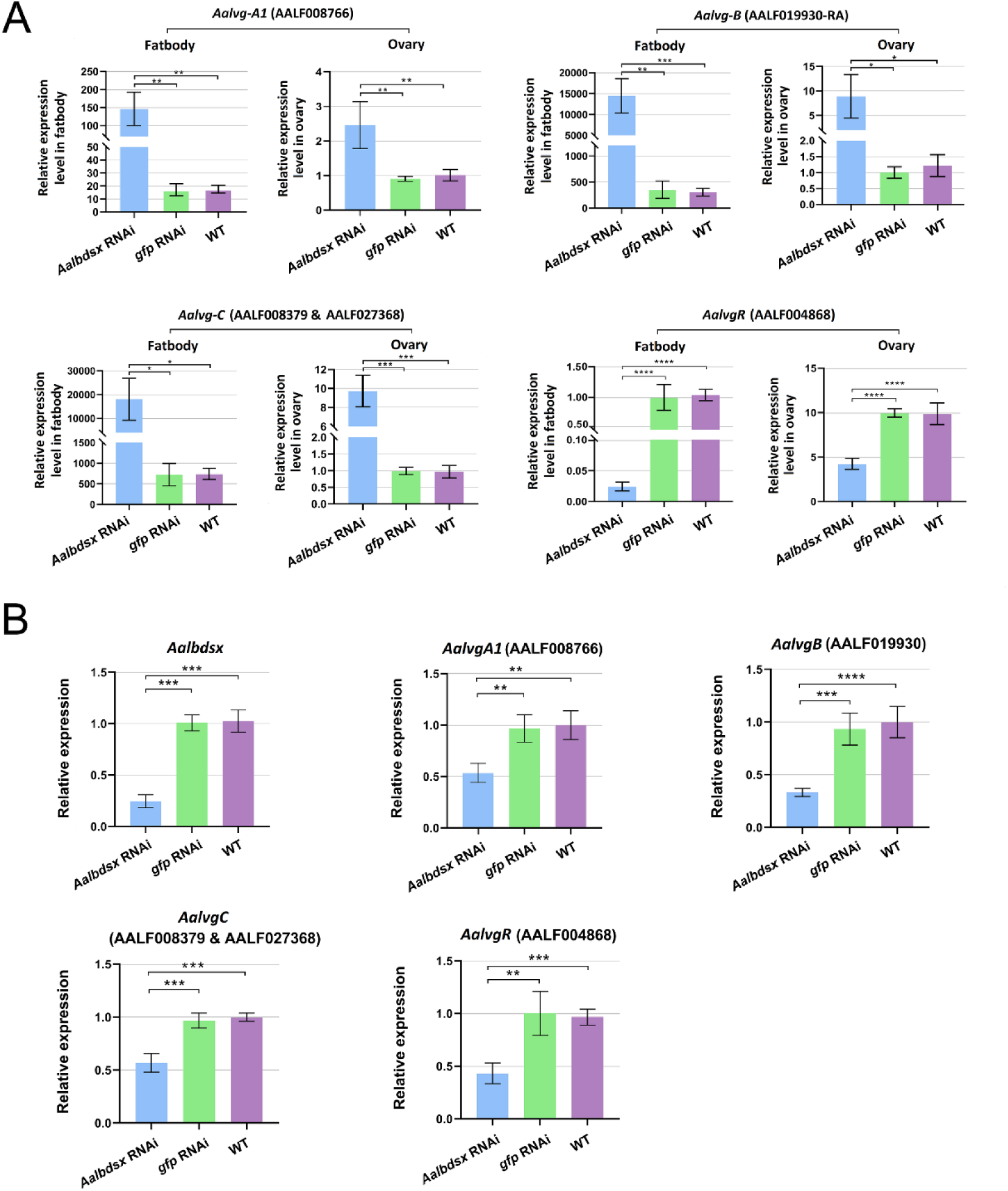
Effect of *Aalbdsx* knockdown on *Vg* and *VgR* gene transcription in *Ae. albopictus*. (A) Effect of *Aalbdsx* knockdown on *Vg* and *VgR* transcription in adult females PBM. (B) Effect of *Aalbdsx* knockdown on *Vg* and *VgR* gene transcription in nonblood-fed adult females. The relative expression of four *Vg* genes and one *VgR* gene was quantified in female mosquitos from the *Aalbdsx* RNAi groups, control groups and WT group. Because of the high nucleotide sequence identity between *AalVg-C1* and *AalVg-C2*, the sum of the transcript levels of the two genes was quantitatively analyzed. The x-axis indicates the groups, and all data are presented as the mean ± SD (unpaired t-test, n = 3; *p < 0.05, **p < 0.01, ***p < 0.005, ****p < 0.001).

### 3.6 High expression of *AalVg* mainly originates from feedback regulation of multiple *AalVg* signaling pathways

The high *AalVg* expression observed in *Aalbdsx*-knockdown blood-fed mosquitoes was obviously contrary to a previous report that DSX is a transcriptional activator of *Vg* in insects(19, 31, 61). Considering the multiple signaling pathways involved in *Vg* and *VgR* regulation post blood feeding(62, 63), we further compared the *Vg* and *VgR* transcript levels of the *Aalbdsx*-knockdown nonblood-fed females with those of the controls. qRT-PCR revealed clearly different results from blood-fed mosquitoes, and *AalVg-A1, AalVg-B, AalVg-C1/C2* and *AalVgR* were all downregulated in the *Aalbdsx* RNAi group (Fig.5B). Our DGE analysis also showed that the *AalVgA1* transcript level was decreased in *Aalbdsx* RNAi mosquitoes. Together, these results indicated that *Aalbdsx* acted as a transcription factor (TF) and activated the transcription of *Vg* and *VgR* in *Ae. albopictus*.

However, the question of why *AalVgs* transcript levels were upregulated in blood-fed females remained. *Vg* is regulated by multiple signaling pathways(62), among which the main cascades include the following: (1) The juvenile hormone (JH) signaling pathway. JH induces the heterodimerization of methoprene-tolerant (Met) with its partner Fushi Tarazu Factor 1 interacting steroid receptor coactivator/steroid receptor coactivator/Taiman (FISC/SRC/Tai) to bind E-box-like motifs in the regulatory regions of *Vg* genes and activate transcription(64-68). (2) The ecdysone signaling cascade. After a blood meal, in the presence of 20 hydroxyecdysone (20E), the ecdysone receptor (EcR) displaces Aedes hormone receptor 38 (AHR38) and forms a heterodimer with the ultraspiracle protein (USP); then, EcR/USP binds to the ecdysone response elements (EcRE) located in the *Vg* promoter and activates expression(69-72). (3) The nutritional amino acid (AA)/ target-of-rapamycin (TOR) signaling pathway. Hemolymph amino acid levels regulate TOR and the transmembrane receptor Frizzled 2 (Fz2), inducing the target gene phosphorylation of S6 kinase (S6K) in the fat body(73, 74). Then, upregulated S6K activates the translation level of the GATA factor, which is a key regulatory factor in *Vg* gene transcription(75, 76). (4) The insulin-like peptide (ILP) signaling pathway. The expression of the ILP in the fat body and brain leads to forkhead box protein O (FOXO) phosphorylation, which in turn triggers *Vg* expression(77-79) (Fig.6A).

**Figure 6.**
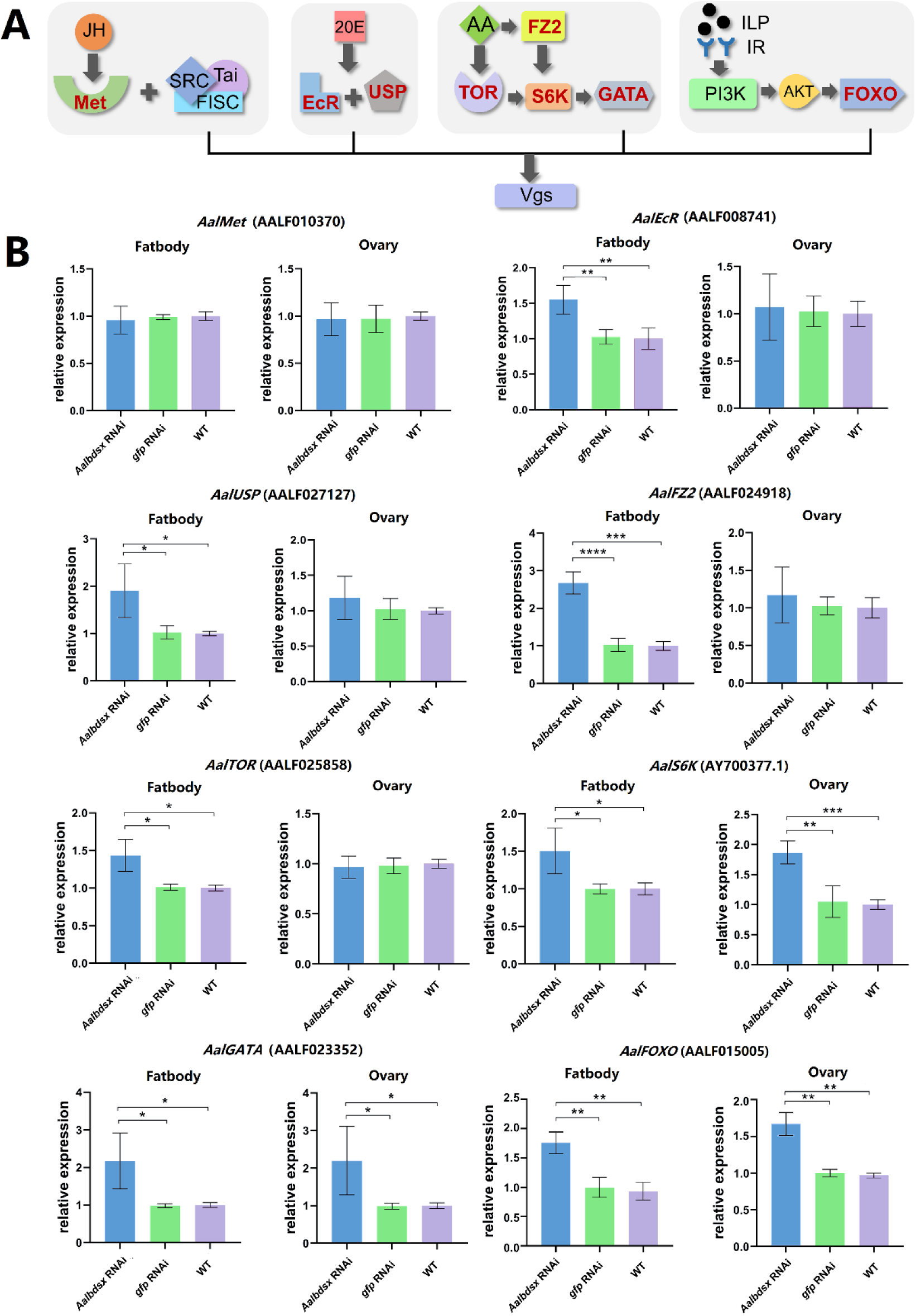
Effect of *Aalbdsx* knockdown on *Vg* pathway-related gene transcription in *Ae. albopictus*. (A) *Vg* is mainly regulated by four signaling pathways. Red font indicates the key genes in these pathways. (B) Effect of *Aalbdsx* knockdown on core genes related to *Vg* pathway transcription in blood-fed adult females. The x-axis indicates the groups, and all data are presented as the mean ± SD (unpaired t-test, n = 3; *p < 0.05, **p < 0.01, ***p < 0.005, ****p < 0.001).

Therefore, we detected the transcript levels of major factors involved in four *Vg* regulatory cascades in *Aalbdsx* RNAi mosquitoes. As Fig.6B shows, in the JH pathway, there was no significant difference in *Met* transcript levels between the *Aalbdsx*-knockdown and control mosquitoes. In the 20E cascades, both *EcR* and *USP* transcript levels were obviously upregulated in the fat body in the *Aalbdsx*-knockdown group compared to the control groups. In the AA signaling pathway, the transcript levels of four genes, *TOR, Fz2, S6K*, and *GATA*, were significantly increased in the fat body under *Aalbdsx*-repressing conditions. *S6K* and *GATA* were also upregulated in the ovary. In the ILP pathway, the *FOXO* level in the fat body was increased in *Aalbdsx* RNAi mosquitoes compared to that in the control mosquitoes (Fig.6B). These results suggested that although *Aalbdsx* acted as a transcriptional activator of both *AalVg* and *AalVgR, Aalbdsx* RNAi caused a decrease in the *VgR* protein content in the ovary and reduced yolk deposition in the ovaries. Intraovarian *Vg* cannot meet the requirement for ovarian development and results in feedback to the fat body to induce *Vg* expression via multiple signaling pathways.

## 4. Discussion

*Dsx* is a central nexus gene at the bottom of the insect somatic sex determination hierarchy that directly or indirectly controls sexually dimorphic morphology, anatomy, physiology, and behavior(80, 81). Direct or indirect Dsx target genes have been described in *D. melanogaster*(28, 29, 40, 82-85), *Tribolium castaneum*(19), *Bombyx mor*i(17, 34, 36, 37, 86), *Bactrocera dorsalis*(13), *Antheraea assama* and *Antheraea mylitta*(17), dung beetles (Onthophagus), rhinoceros beetles (Trypoxylus)(18, 87), and *Cyclommatus metallifer*(88). In some insects, such as *Nasonia vitripennis*, O*nthophagus taurus, Onthophagus sagittarius, Trypoxylus dichotomus*, and *Papilio polytes*, although the target genes of *dsx* have not been determined, obvious morphological changes are observed post-*dsx* repression(89).

The effect of *dsx* repression on mosquitoes has also been demonstrated in *An. gambiae* and *Ae. aegypti*. The knockdown of female-specific *dsx* in *An. gambiae* larvae can cause a decrease in pupation and emergence rates, shortened lifespans and reduced fecundity of females(90, 91). *Dsx* silencing in *Ae. aegypti* pupae leads to decreases in wing size, proboscis length and the lifespan of adult females, in addition to decreased ovariole length and a reduced ovariole number in blood-fed adult females. Moreover, *Dsx* downregulation can reduce the length of the female antennae and maxillary palps and sensilla distributed on their surface(45). Although *dsx* repression disrupts multiple sex-related morphological, physiological, and behavioral traits of adult females, the target gene repertoire regulated by *dsx* in the mosquito is entirely unknown, except that several *odorant receptor* (*OR*) genes have been shown to be affected by *dsx* interference(45).

In this study, we first screened differentially expressed genes in a DGE library post-*dsx* repression and then confirmed direct candidate target genes of *dsx* by qPCR and ChIP-qPCR. We used total RNA extracted from mosquito whole bodies to construct the libraries rather than tissues that are dimorphic between the sexes, which might have caused some tissue-biased and low-abundance differentially expressed transcripts to be neglected, ultimately leading to the screening of relatively few differentially expressed genes from our libraries. As a result, the most downregulated gene was *vitellogenin-A1 precursor* (*VgA1*), a direct target of *dsx* in a variety of insects. Vitellogenesis (process of yolk formation in the oocyte) is a key physiological event in mosquito life and is generally dependent on the availability of a protein-rich blood meal to provide nutrition for the developing embryo(92). The Knockdown of the *VgA1*(also known as *Vg-2a*) gene via RNAi restores host-seeking behavior in young *Ae. albopictus* females(93). The second most significantly downregulated gene was *elongation factor-1α* (*EF-1α*) (AALF007667(59)). *EF-1α* promotes the delivery of aminoacyl-Trna which shows GTP-dependent binding to the acceptor site of the ribosome during protein biosynthesis. *EF-1α* plays important roles in insect development and reproduction. *EF-1α* RNAi increases nymph mortality and severely hinders the ovarian development of the fall armyworm *Spodoptera frugiperda*(94); this factor also affects the lifespan of *D*.*melanogaster*(95). The third most significantly downregulated gene was *allantoinase* (*ALN*). *ALN* catalyzes the hydrolysis of allantoin to produce allantoic acid, which is then imported in insects(96), *ALN* is involved in the metabolic pathways of uric acid degradation and urea synthesis in adult female *Ae. aegyp*ti, shows relatively high expression in the fat body and Malpighian tubules and is upregulated after a blood meal, which guarantees the survival of blood-fed mosquitoes(97). The *myosin V* gene (*MyoV*) (AALF004459 (59) was also among the top ten downregulated genes. The RNAi knockdown of myosin family genes can reduce wing and leg size, inhibit eclosion and decrease fertility and egg hatchability in *T*.*castaneum*(98). *MyoV* mutants show a high frequency of larval lethality, which is also observed in *Drosophila*, indicating that *MyoV* is an essential gene (99). Furthermore, seven of the target genes are putative salivary secreted proteins or predicted salivary secreted peptides/proteins. Salivary secreted proteins are critical for successful blood feeding because they facilitate blood vessel location and blood ingestion in female mosquitoes(100, 101) and may function as lubricants of the mouthparts(102), which indicates that *dsx* might be involved in blood feeding and ingestion processes. The odorant-binding protein OBP23 was significantly upregulated post-*dsx* disruption. *OBP23* is a typical gene showing male-biased high expression in *Ae. aegypti*(103), which suggests that female *dsx* negatively regulates the expression of some male-biased genes in females.

Although *dsx* sequences are relatively conserved, the target genes of *dsx* evolve rapidly in response to insect sexual dimorphism(104), and to accommodate evolutionarily novel traits(30, 83). However, among these target genes, *Vg* is relatively conserved in insects and has been functionally validated as a direct *dsx* target(13, 17, 19, 29, 31-35).

In mosquitoes, massive amounts of YPPs are produced exclusively in the fat body after a blood meal, which are then released in vitellogenic females and transported to developing oocytes by receptor-mediated endocytosis(105). *Vg* expression controls extremely complex gene regulatory networks. In addition, four major cascades (the JH, 20E AAs, and ILP signaling pathways) and a number of miRNAs are also involved in the *Vg* regulation cascade in mosquitoes(106-108). Our qPCR and ChIP-qPCR results confirmed that the four *Vg* genes were direct genes of *dsx* in *Ae. albopictus*; furthermore, *dsx*-knockdown led to the repression of all *AalVgs* in adult females, and the transcript levels of the *AalVgs* showed an obvious increase induced by a blood meal. Moreover, the transcript levels of major regulatory factors involved in four major *Vg* signaling pathways were typically increased post-blood meal. These results indicated that compared to the downregulation caused by *dsx* repression, induction effects effect on four major cascades involved in *Vg* regulation predominate post-blood meal.

*VgR* belongs to the low-density lipoprotein receptor (LDLR) gene superfamily(105) and is primarily detected in insect ovarian tissue, primarily in the oocytes(109-111). In *Colaphellus bowringi, VgR* gene transcription is regulated by JH via the intracellular JH receptor *Met* and the JH-responsive transcription factor *Krüppel homolog 1* (*Kr-h1*) (112). Ecdysone-responsive early gene-encoded protein-binding sites were also found in the *Ae. aegypti VgR* promoter region(111). Our results confirmed that *VgR* was a direct target gene of *dsx*. Furthermore, *dsx* interference led to the repression of all *AalVgR* genes in both non-blood-meal-fed and blood-meal-fed adult females, which suggests that *dsx* plays a main regulatory role in the competition of *AalVgR* with other regulatory pathways post-blood meal, which ultimately leads to the repression of ovarian development. However, the possibility that other unknown DSX target genes lead to ovarian defects and result in the downregulation of *VgR* transcripts should not be excluded.

Furthermore, we cloned and characterized *dsx* in *Ae. albopictus*, a secondary dengue vector in Asia. The direct target genes of *Aalbdsx* were confirmed by DGE combined with ChIP-qPCR analysis, which provided molecular evidence supporting possible explanations for the disruption of dimorphic morphology, anatomy, physiology, and behavior described in *dsx*-silenced females in previous work. The conservation of *Vg* and *VgR* as direct *dsx* target genes observed in other insects was confirmed in mosquitoes; however, the effects of *Aalbdsx* RNAi on the ovarian development of adult females post-blood meal were mainly exerted via *AalVgR*, rather than *AalVgs*. The results of our study not only provide a reference for the further elucidation of the sex determination cascade and comparative analyses of *dsx* target interactions in mosquitoes but also reveal potential molecular targets for application to the development of sterile male mosquitoes to be released for vector control.

## Acknowledgements

The authors thank Prof. Zhijian Tu at Virginia Polytechnic Institute and State University for his valuable suggestions regarding the manuscript. We are grateful to BGI (Beijing Genomics Institute) for sequencing technical advice. This study was funded by the National Natural Science Foundation of China (81672054, 81871688 and 31830087), the National Institutes of Health, USA (AI136850), and the Natural Science Foundation of Guangdong Province (2017A030313120).

